# A comprehensive stroke risk assessment by combining atrial computational fluid dynamics simulations and functional patient data

**DOI:** 10.1101/2024.01.11.575156

**Authors:** Alberto Zingaro, Zan Ahmad, Eugene Kholmovski, Kensuke Sakata, Luca Dede’, Alan K. Morris, Alfio Quarteroni, Natalia A. Trayanova

## Abstract

Stroke, a major global health concern often rooted in cardiac dynamics, demands precise risk evaluation for targeted intervention. Current risk models, like the CHA_2_DS_2_-VASc score, often lack the granularity required for personalized predictions. In this study, we present a nuanced and thorough stroke risk assessment by integrating functional insights from cardiac magnetic resonance (CMR) with patient-specific computational fluid dynamics (CFD) simulations. Our cohort, evenly split between control and stroke groups, comprises eight patients. Utilizing CINE CMR, we compute kinematic features, revealing smaller left atrial volumes for stroke patients. The incorporation of patient-specific atrial displacement into our hemodynamic simulations unveils the influence of atrial compliance on the flow fields, emphasizing the importance of LA motion in CFD simulations and challenging the conventional rigid wall assumption in hemodynamics models. Standardizing hemodynamic features with functional metrics enhances the differentiation between stroke and control cases. While standalone assessments provide limited clarity, the synergistic fusion of CMR-derived functional data and patient-informed CFD simulations offers a personalized and mechanistic understanding, distinctly segregating stroke from control cases. Specifically, our investigation reveals a crucial clinical insight: normalizing hemodynamic features based on ejection fraction fails to differentiate between stroke and control patients. Differently, when normalized with stroke volume, a clear and clinically significant distinction emerges and this holds true for both the left atrium and its appendage, providing valuable implications for precise stroke risk assessment in clinical settings. This work introduces a novel framework for seamlessly integrating hemodynamic and functional metrics, laying the groundwork for improved predictive models, and highlighting the significance of motion-informed, personalized risk assessments.

## Introduction

Stroke remains a formidable medical challenge worldwide, posing a significant threat to both life and quality of life. According to the World Stroke Organization, 13 million people experience stroke per year and, of those, 5.5 million die as a result^1^. Characterized by a sudden interruption of blood flow to a part of the brain, stroke often results in long-lasting neurological impairments^2^. The left atrium (LA) and left atrial appendage (LAA) are known to be pivotal in the multifaceted etiology of stroke^3^. Thrombi, or clots, in this chamber, once formed, can embolize and travel to the cerebral circulation, precipitating an ischemic stroke. Anomalies in the LA/LAA shape and function have been increasingly recognized as a potential precursor of cerebrovascular events such as ischemic stroke^4,5^, necessitating a closer examination of the intricate morphological, functional, and hemodynamic characteristics of this cardiac chamber. Atrial fibrillation (AF), a common arrhythmia, is often associated with alterations in LA structure and biomechanics, contributing to stasis of blood flow and fostering a pro-thrombotic state within the atrium^6,7^. Congestive heart failure and hypertension are also linked with increased stroke risk^8–10^. Thus, it is clear that a deeper understanding of LA hemodynamics can pave the way to the development of predictive models for stroke risk in susceptible populations.

Contemporary risk assessment tools, such as the CHA_2_DS_2_-VASc score, while ubiquitously used, predominantly focus on patient-centric risk factors, such as age, sex, and comorbidities, and do not consider factors pertaining to LA morphology and function^11^. Oral anticoagulant (OAC) therapy is prescribed for AF patients based on this score^12^. However, OACs have the potential to cause severe internal bleeding including intracerebral hemorrhage and are associated with higher mortality rates^11,13^, necessitating an accurate assessment of OAC need. Unfortunately, the CHA_2_DS_2_-VASc score has severe limitations in its predictive capabilities of stroke. This *one-size-fits-all* approach has been shown to be inaccurate and inconsistent over various patient cohorts^14,15^. Several studies showed that patients might be administered too many OACs and exposed to its side effects without deriving any benefit from it, or are incorrectly classified as low risk and thus are not protected^11,16,17^. In addition, for patients who do not have AF, or in patients that exhibit asymptomatic arrhythmias and atrial high-rate episodes, it is difficult to accurately assess stroke risk, since the CHA_2_DS_2_-VASc is not directly applicable^18,19^. Therefore, it is crucial to precisely identify the patients who truly require these treatments, ensuring that high-risk individuals are not misclassified as low risk, while also avoiding unnecessary exposure to the side effects of the medications and procedures administered for those at lower risk. This gap in current risk stratification models underscores an urgent need for a more nuanced, individualized approach that encapsulates the hemodynamic intricacies and geometric variabilities inherent within the LA. This need can be addressed by the use of in-silico methodologies as a noninvasive tool to predict the likelihood of thrombus formation and stroke risk in an individual subject^20–32^.

The use of computational fluid dynamics (CFD) models to assess hemodynamics in the LA/LAA has gained significant attention in recent years for the purposes of thrombosis and stroke risk assessment, specifically in cohorts of AF patients. For this purpose, Garcia et al. used rigid wall assumptions for CFD simulations on idealized ellipsoidal geometries synthetically created to resemble the atrium^33^. Many studies utilized patient-specific 3D computed tomography (CT) images to simulate flow dynamics across distinct LAA morphologies in AF and stroke patients. However, the majority of these studies ignore the contractility of the atrial chamber by carrying out simulations under the assumption of a rigid LA wall ^34–40^. In contrast, recent studies have demonstrated that considering the contractility motion of the atrial walls, as opposed to rigid assumptions, yields distinct flow fields in the atrium^22,26,27,41^. As a matter of fact, the rigid wall assumption tends to overestimate thrombosis risk^37^ and is representative of a very specific and extreme case of AF (chronic AF)^42^. In order to account for endocardial motion in CFD simulations, some researchers have developed methods for artificially computing the wall LA displacement to study stroke risk stratification. Musotto et al.’s work utilized idealized assumptions for LAA mechanical properties in CFD simulations^43^. Masci et al. introduced a definition of wall motion by applying a random displacement function to AF patient-specific 3D geometries^44^. Zingaro et al. leveraged an idealized displacement model with a 0D closed-loop circulation model to compute an LA boundary motion^45^. The method was then extended by Corti et al. to study the impact of AF on hemodynamics^21^. However, in the absence of clinical data, all these displacements remained idealized and non-patient-specific. Differently, with the use of dynamic patient-specific data such as CINE MRI or 4D CT scans, hemodynamic models can be more realistic and patient-specific. Few examples of patient-specific displacement-informed CFD simulations are available in the literature; however, they all focus on a small cohort of patients. Garcia-Villalba et al. perform CFD simulations in a cohort of six patients and use 4D CT scans for boundary displacement in AF patients with and without thrombus formation^26^. Otani et al. carried out CFD simulations with displacement from CINE MRI in two AF patients^46^. A 3D interpolation algorithm was applied by Qureshi et al. for smoothing and minimization of discontinuities between slices to form a single patient-specific surface mesh of the LA with CINE MRI^39,47^. Dueñas-Pamplona et al. used dynamic 4D CT data to simulate an LA CFD model in two AF patients for the purpose of model verification and proof of concept of using patient-specific boundary conditions. The insufficiency of these approaches – both in terms of wall modeling and size of the patient cohort – highlights the urgent need for more comprehensive studies accounting for atrial contractility and a larger number of patients to achieve a more personalized and mechanistic assessment of stroke risk.

In this paper, we assess stroke risk through atrial CFD simulations, incorporating patient-specific wall displacement derived from image data. We consider a cohort of eight patients, evenly divided into stroke and control cases. We do not restrict the patient cohort to AF patients as many of the aforementioned works do, allowing for generalizable insights into the correlation between LA hemodynamics and stroke, independently of the precurring pathology. To run personalized hemodynamic simulations, we merge CINE cardiac magnetic resonance (CMR) and contrast-enhanced (CE) magnetic resonance angiography (MRA) to obtain accurate time-deforming volumetric meshes. To begin, we analyzed functional and morphological characteristics of the LA and LAA directly from the CMR to identify distinguishing biomarkers between the two patient groups. Then, we performed CFD simulations driven by the patient-specific displacements. We found that hemodynamic features from the LA chamber alone do not separate well between the stroke and control groups of our cohort. Differently, when we plot our hemodynamic computations against the functional and morphological features, we find several fluid dynamics features to be in a separate range for the stroke group. Specifically, we provide a comprehensive merging of all our computed hemodynamic features with the functional data and morphological characteristics of the LA and LAA. We report that several of these new quantities provide clear distinctions between the stroke patients and control patients in our cohort, especially when normalized with maximum volume, stroke volume, and beats per minute. This study presents an innovative framework that seamlessly integrates hemodynamic and functional metrics, establishing a foundation for enhanced predictive models. It underscores the vital role of motion-informed, personalized risk assessments. Importantly, no other study has explored the integration of hemodynamic features with functional or morphological computations to achieve a personalized and comprehensive stroke risk assessment.

## Results

To unravel the biomarkers associated with the risk of stroke development, we systematically compute and analyze various features related to the LA/LAA geometry, its functioning, and hemodynamics. Starting with patient-specific data, we introduce a set of biomarkers encoding information on the geometry and functioning of the LA and its appendage. Subsequently, our focus shifts to the hemodynamics of both the LA and LAA, exploring the outcomes derived from the personalized CFD simulations. In conclusion, in a cohesive approach aimed at providing a more personalized and mechanistic understanding of each patient, we integrate these insights by combining functional data with fluid dynamics features.

### Functional data

Our patient cohort consists of eight patients: four control, and four stroke cases. Patient characteristics are shown in Table 1. Stroke patients presented higher beats per minute (BPM), hence a lower heartbeat period. To assess geometry and functioning of the LA, in Figure 1, we present a set of biomarkers derived from dynamic meshes, which were generated based on CMR data. Notably, control patients, with the exception of C1, show larger maximum and minimum LA volumes in comparison to stroke patients. Moreover, stroke patients generally exhibit lower stroke volumes (SV) compared to the control group. However, in terms of ejection fraction (EF) of the LA chamber, a clear differentiation between the two categories is not readily apparent. Additionally, we perform a similar analysis on the LAA across various time frames. Similar to the LA, we observe that control cases tend to have a larger LAA volume, with the exception of C1. Furthermore, similar to the primary chamber, stroke patients typically exhibit reduced SV, while EF does not distinguish between the two categories. For the LAA, we additionally compute the ostium area (OA) and the tortuosity (*τ*) at the time corresponding to the minimum LA volume, which coincides with the closure of the mitral valve (MV). Large tortuosity values usually reflect the complexity of LAA shape^44^. Nonetheless, both of these features did not reveal specific discernible patterns. It is worth noting that stroke patients generally demonstrated a narrower range of variation in these measurements when compared to the control group, despite the lack of a distinctive pattern.

**Table 1.** Information of the eight patients considered in the study.

|  | patient ID | sex | age | other pathologies | BPM | Period [s] |
| --- | --- | --- | --- | --- | --- | --- |
| **Control** | C1 | M | 44 | none | 57.1 | 1.05 |
|  | C2 | M | 56 | none | 50.3 | 1.19 |
|  | C3 | M | 45 | none | 57.7 | 1.04 |
|  | C4 | M | 49 | none | 51.5 | 1.17 |
| **Stroke** | S1 | F | 29 | none | 76 | 0.79 |
|  | S2 | M | 65 | hypertension | 67 | 0.90 |
|  | S3 | F | 69 | hypertension | 51.5 | 1.17 |
|  | S4 | F | 60 | diabetes mellitus | 69 | 0.87 |

**Figure 1.**
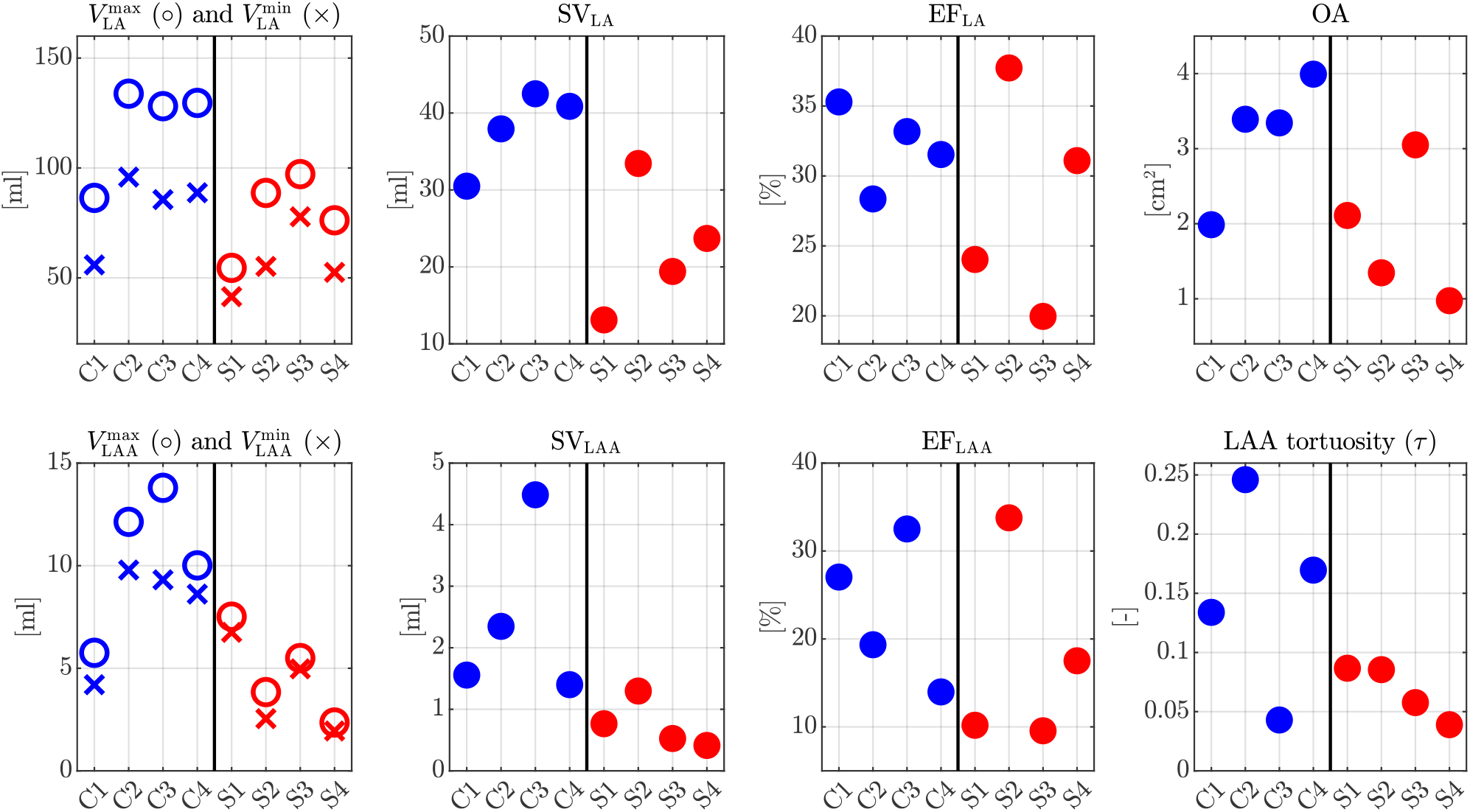
Biomarkers computed from dynamic meshes. In blue 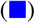, control cases; in red 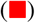, stroke cases. From the top left to bottom right: maximum (circle) and minimum (cross) LA volume, LA stroke volume, LA ejection fraction, ostium area of LAA, maximum (circle) and minimum (circle) LAA volume, LAA stroke volume, LAA ejection fraction, tortuosity of the LAA.

### Results from hemodynamic simulations

To explore the atrial hemodynamics, we now showcase the outcomes of our personalized CFD simulations. Figure 2 presents the transient data of LA volumes and flowrates calculated at the MV section for both control and stroke cases. Consistent with our functional data, stroke patients consistently exhibit diminished LA volumes. Similarly, our simulation results indicate reduced flowrates at the valve section in stroke patients. Additionally, we evaluate the MV flowrate during the E-wave and A-wave for all patients, as detailed in Table 2. Notably, our findings reveal that the EA ratio does not serve as a discriminating factor for assessing stroke risk.

**Table 2.** MV flowrate peaks at E-wave and A-wave; EA ratio.

|  |  | Control |  |  |  | Stroke |  |  |  |
| --- | --- | --- | --- | --- | --- | --- | --- | --- | --- |
|  |  | C1 | C2 | C3 | C4 | S1 | S2 | S3 | S4 |
| **E-wave** | [mL/s] | 433.0 | 799.8 | 568.7 | 717.0 | 430.8 | 432.1 | 484.7 | 475.5 |
| **A-wave** | [mL/s] | 336.9 | 817.4 | 512.9 | 632.7 | 298.1 | 492.5 | 469.4 | 401.1 |
| **EA ratio** | [-] | 1.29 | 0.98 | 1.11 | 1.13 | 1.45 | 0.88 | 1.03 | 1.18 |

**Figure 2.**
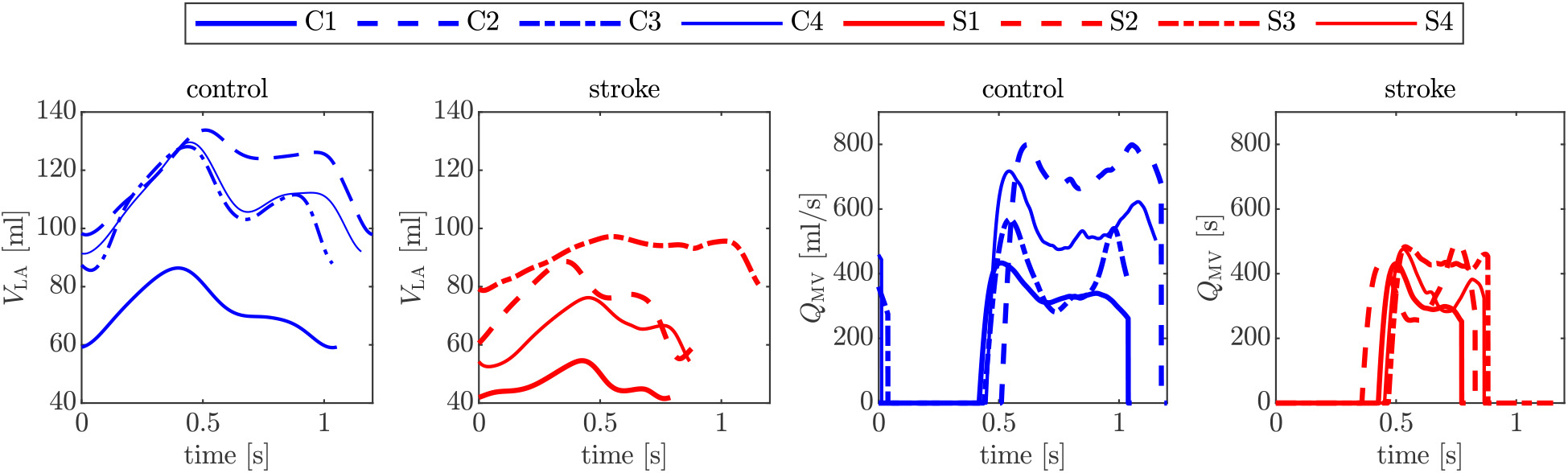
LA volumes and MV flowrate versus time for the control and stroke cases. In blue 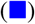, control cases; in red 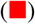, stroke cases.

Figure 3 is a visual representation of the phase-averaged blood velocity captured at the A-wave peaks. These time points were determined by examining the transient MV flowrates (Figure 2). A common observation across all cases reveals a dynamic mixing of blood within the LA, as it courses toward the MV section, giving rise to an intricate blood flow pattern. Our analysis indicates larger velocities in proximity to the pulmonary veins, while the blood flow in the LAA exhibits a considerably slower pace. In stark contrast to the control cases, stroke patients consistently display reduced velocity in the pulmonary veins, the LA, and its appendage. This aligns with the MV flowrate measurements depicted in Figure 2.

**Figure 3.**
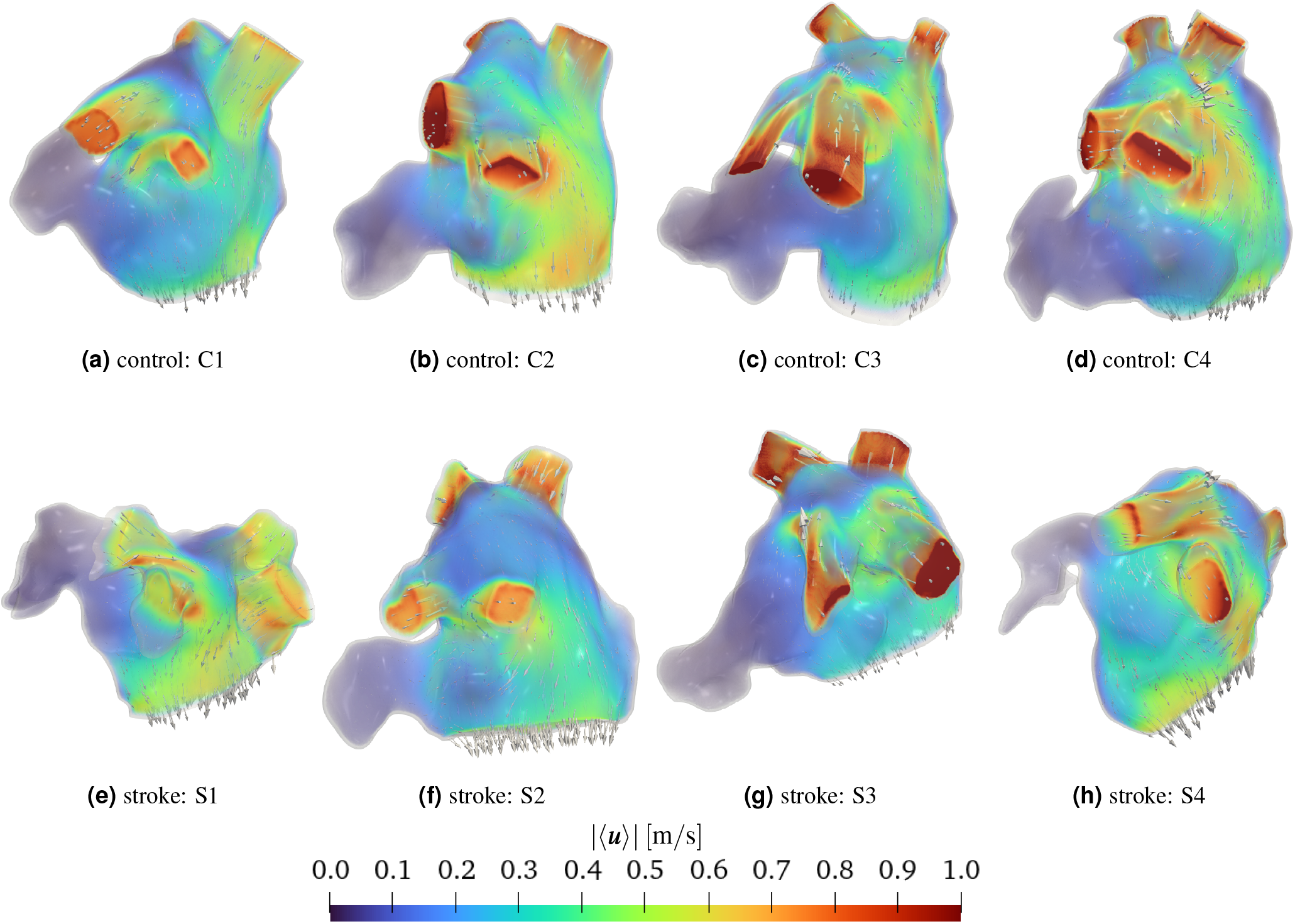
Volume rendering of phase-averaged velocity magnitude with glyphs of velocity at the A-wave peak. Top (a-d): control cases; bottom (e-h): stroke cases.

To assess comprehensively the impact of stroke on atrial flows, we calculate various hemodynamic features that gauge endothelial susceptibility and the likelihood of thrombus formation. These include:

- Flow Stasis (FS), representing the fraction of time during a heartbeat in which the velocity falls below 0.1 m*/*s^21,48^.
- Time Averaged Wall Shear Stress (TAWSS), serving as an average measure of shear stress at endocardial walls.
- Oscillatory Shear Index (OSI), quantifying the degree of change in wall shear stress (WSS) throughout the cardiac cycle.
- Relative Residence Time (RRT), proportional to the residence time of blood particles near the wall, identifying areas with both low and oscillatory WSS^49,50^.
- Endothelial Cell Activation Potential (ECAP), detecting regions characterized by oscillatory and low-stress conditions^51^.

In Figure 4, boxplots illustrate the distribution of hemodynamic features computed for the entire LA at the top and its auricle at the bottom, discerning between stroke and non-stroke cases. Notably, both groups display substantial variability in results, marked also by the presence of numerous outliers. Focusing on the median values, our analysis for the LA reveals that all the examined features effectively differentiate between stroke and non-stroke cases, with the exception of S4 (see also Supplementary Table 1 in Supplementary Material). Specifically, stroke cases exhibit higher FS, reduced TAWSS, and elevated OSI, RRT and ECAP. This pattern is notably pronounced in the LA. In contrast, the LAA does not exhibit a discernible distinction between the two subgroups (results for the LAA are provided in Supplementary Figure 1 and Supplementary Table 1 in Supplementary Material). The nuanced variations between the main chamber and its auricle underscore the complexity of hemodynamic responses in stroke and non-stroke scenarios. This also suggests the need to incorporate additional patient-specific data into our analysis, such as anatomical and functional measurements from CMR scans.

**Figure 4.**
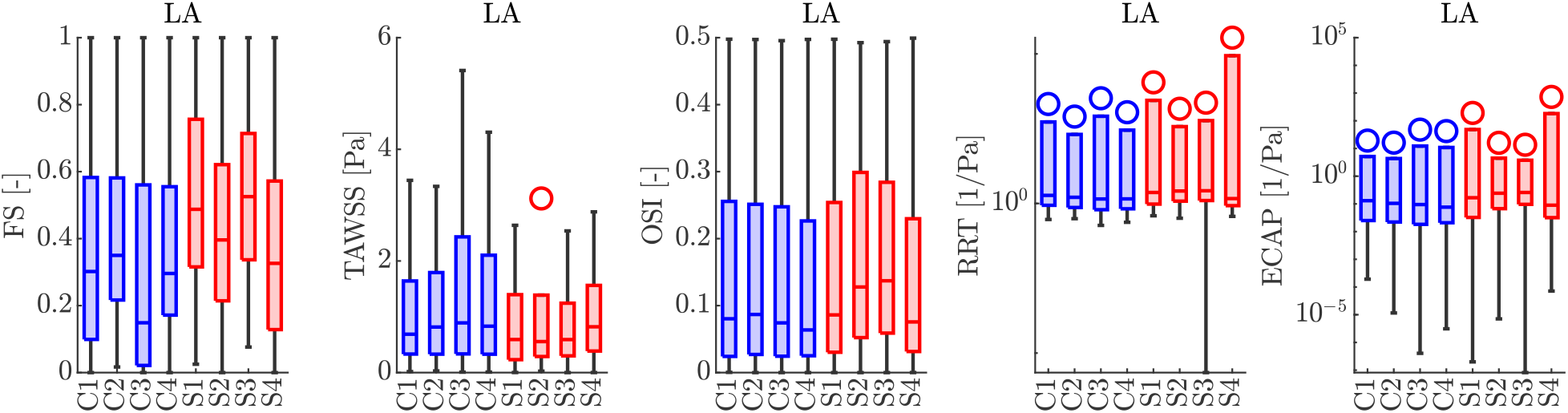
Boxplots of hemodynamics features computed with patient-specific CFD simulations in the LA. From the left to the right: FS, TAWSS, OSI, RRT, and ECAP. In blue 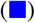, control cases; in red 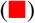, stroke cases. The same analysis for the LAA is provided in Supplementary Figure 1 in Supplementary Material.

### Merging functional data with hemodynamics features

In our pursuit of a comprehensive understanding of atrial hemodynamics and its implications for predicting stroke risk, we focus on the median values of each hemodynamic feature. We plot these values against a set of biomarkers directly derived from CMR data, encompassing functional parameters such as maximum and minimum LA volumes, SV, EF, and BPM. We show these results in Figure 5, leveraging a logistic regression model with a binomial distribution to distinguish between stroke and non-stroke subgroups. The black dashed lines in the plot represent the predictions of the logistic model within the parameter space of interest. The logistic regression line on the plot serves as a predictive boundary. When this line effectively separates the data points (stroke and control groups), it indicates a correlation between the hemodynamic features and functional data in terms of stroke prediction. Remarkably, the maximum LA volume emerges as a robust discriminator between the two groups when plotted against all hemodynamic features, with the exception of TAWSS. Conversely, the minimum volume serves as a discriminant for the FS only. Furthermore, atrial SV consistently proves to be a reliable metric for segregating the subgroups, while the EF of the LA falls short in discerning differences across all hemodynamic features. Interestingly, BPM emerges as a clear distinguishing factor between the two subgroups. Notably, the inclusion of functional measurements, such as SV and BPM, has a transformative effect, reclassifying case S4 from an outlier to a position within the stroke area of the plot.

**Figure 5.**
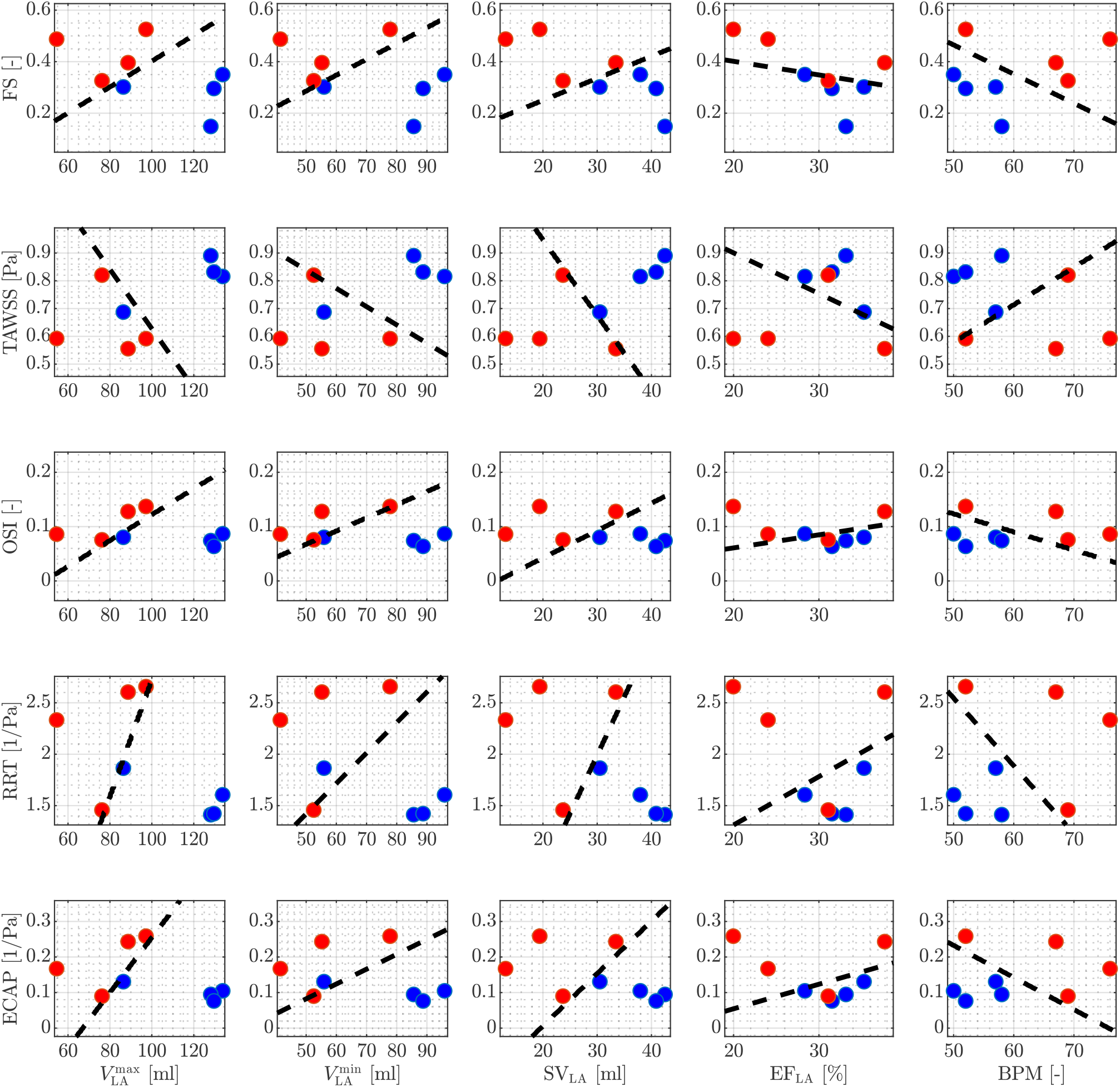
Medians of hemodynamics features in the LA against functional data from CMR. In blue 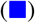, control cases; in red 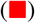, stroke cases. The black-dashed line is obtained by running logistic regression. From top to the bottom: FS, TAWSS, OSI, RRT, and ECAP are plotted against maximum LA volume, minimum LA volume, LA SV, LA EF, and BPM (from left to right).

In Figure 6, we extend our study to focus on the LAA, considering both functional and anatomical measurements. These encompass maximum and minimum LAA volumes, LAA SV, LAA EF, BPM, tortuosity, and OA. In contrast to the LA, identifying a functional feature capable of effectively distinguishing between the stroke and non-stroke subgroups proves to be a more intricate challenge for the LAA. Remarkably, our investigation reveals that LAA SV emerges as a the only robust measure that adeptly characterizes the two subgroups across all considered hemodynamic features in this study. It is noteworthy to mention that, in the absence of any functional measure, the mechanistic insights provided solely by hemodynamic features fall short in distinguishing the two groups for the LAA.

**Figure 6.**
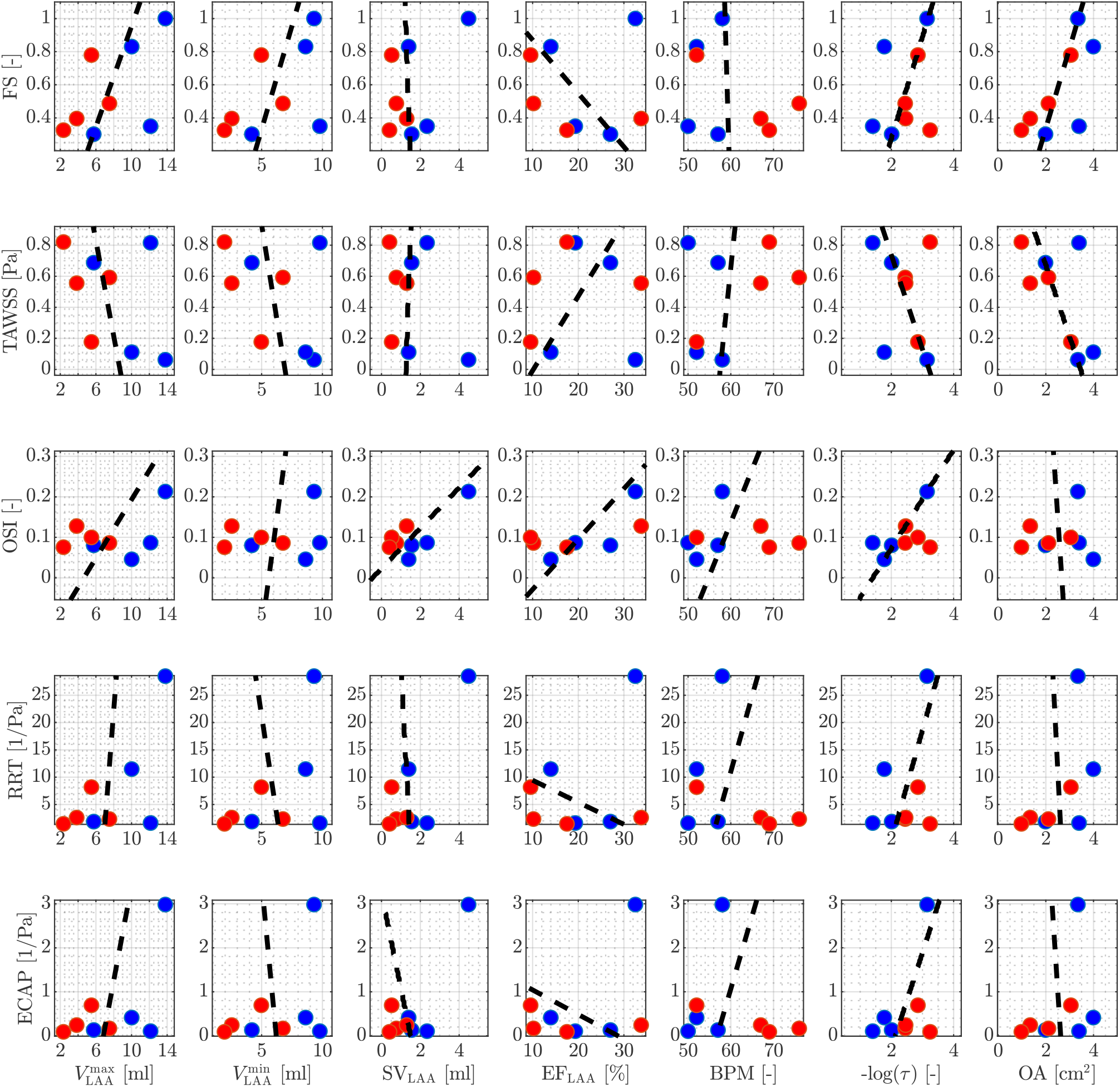
Medians of hemodynamics features in the LAA against functional data from CMR. In blue 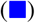, control cases; in red 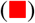, stroke cases. The black-dashed line is obtained by running logistic regression. From top to the bottom: FS, TAWSS, OSI, RRT, and ECAP are plotted against maximum LAA volume, minimum LAA volume, LAA SV, LAA EF, BPM, tortuosity, and OA (from left to right).

### A look into the transitional-to-turbulence regime

To delve into the impact of stroke on the transitional-to-turbulence regime that may occur in atrial flows^50^, in Figure 7a, we present the temporal evolution of kinetic energy (*E*_*k*_) and enstrophy (*S*) for both control and stroke patients. Consistently with elevated flow stasis and diminished velocities, stroke patients exhibit lower kinetic energies. However, the enstrophy, representing vorticity production and dissipation, i.e. a measure of turbulence^50,52,53^, does not reveal a discernible pattern between stroke and control patients.

**Figure 7.**
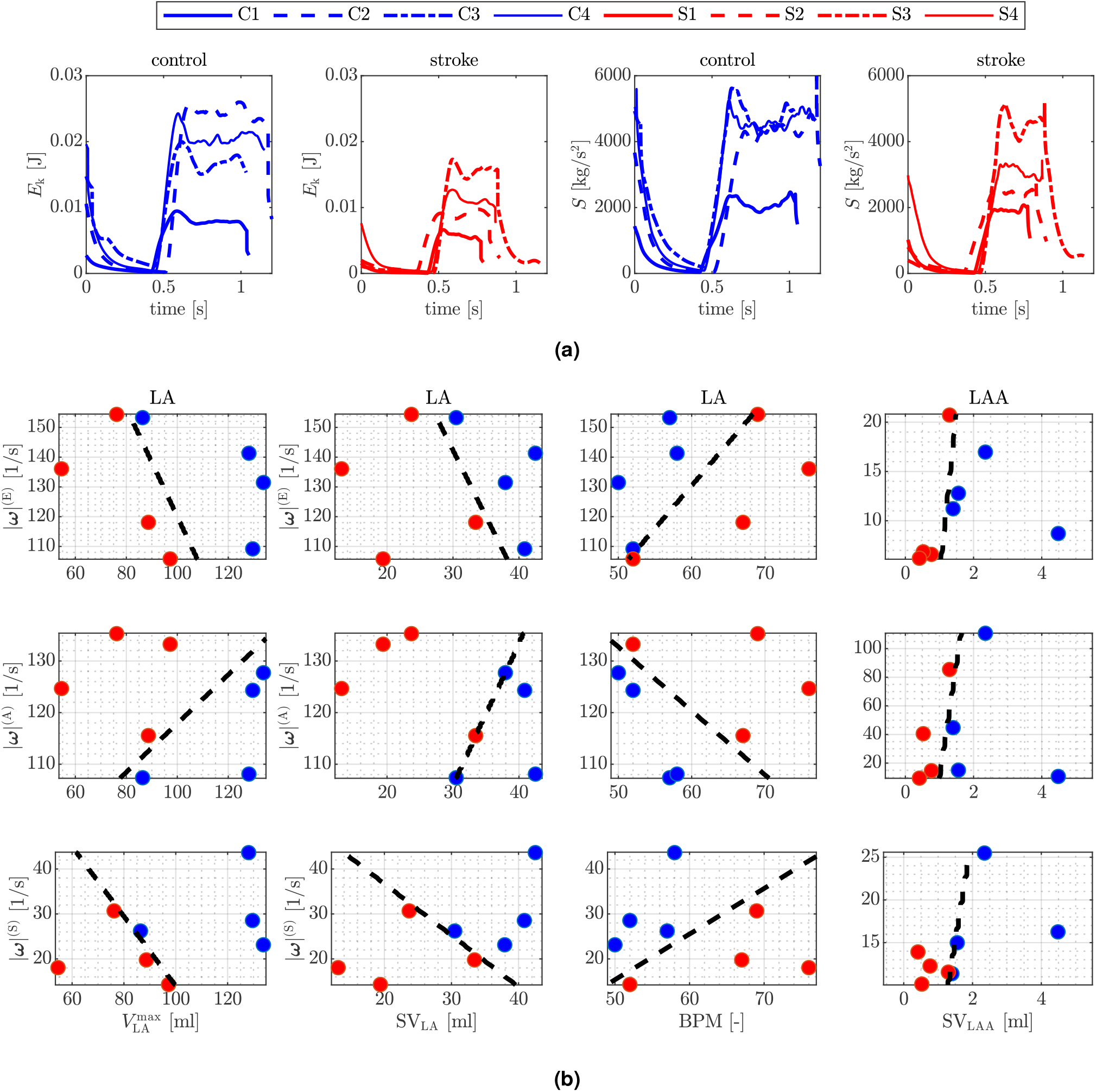
A look into turbulence. (a) Kinetic energy *E*_k_ and enstrophy *S* versus time for the control and stroke cases. (b) Medians of vorticity magnitude against functional data from CMR. In blue 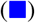, control cases; in red 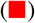, stroke cases. The black-dashed line is obtained by running logistic regression. From top to the bottom: vorticity magnitude at E-wave, A-wave, and systolic peak (MV closed) are plotted against LA maximum volume, LA SV, BPM, and LAA SV (from left to right).

For a more in-depth exploration of the effects of stroke on turbulence, we analyze the vorticity magnitude (|***ω***|) at three distinct phases of the heartbeat: E-wave peak, A-wave peak, and systolic filling (when the MV is closed) for both the LA and the LAA. As with previous features, we plot these results against various functional measurements of the LA. Figure 7b specifically showcases the outcomes for measurements demonstrating correlation. Similar to previous hemodynamic features, we observe correlations with LA maximum volume, LA SV (except for |***ω***|^(A)^, which presents an outlier), BPM, and LAA SV. For a comprehensive view of the additional results against all functional features see Supplementary Figure 2 and Supplementary Figure 3 in Supplementary Material.

## Discussion

The objective of this paper was to deliver precise and personalized assessments of stroke risk through patient-specific LA hemodynamic simulations. The study integrated functional characteristics obtained from CMR with hemodynamic insights derived from patient-specific CFD simulations, which considered the motion of the LA. Standalone analyses of functional and hemodynamic features did not distinctly differentiate between stroke and control cases. However, through a synergistic merging of these variables, a clear distinction between the two groups emerged, offering a personalized and mechanistic assessment of stroke risk.

Our cohort comprised eight patients, evenly split between control and stroke cases. While a majority of previous works on assessing left atrial hemodynamics for stroke and thrombosis risk focused on patients with AF, our cohort does not include any AF patients. This study presents a framework that gives insight into stroke risk metrics for populations who had stroke but who not necessarily had episodes of AF. Indeed, many of the previous computational and clinical studies are solely focused on the link between atrial hemodynamics and stroke risk in AF patients only. This enabled us to provide broader observations regarding the connection between LA hemodynamics, LA functioning and stroke, irrespective of the precurring pathology. Most of the CFD research works on this topic assume rigid LA wall, whereas the works in which a realistic displacement is included are always confined to a small number of patients. In this paper, we incorporated CINE CMR in our patient-specific simulations and, to the best of our knowledge, this is the study with the largest cohort of patients utilizing patient-specific LA motion. This is a crucial aspect in atrial CFD simulations since, as often evidenced in the literature^22,26,27,41^, incorporating motion into hemodynamic simulations yields diverse flow fields in the left atrium, potentially influencing the accuracy of hemodynamic biomarkers used in stroke risk assessment.

By analyzing features directly extracted from CMR, we observed that stroke patients generally exhibited lower volumes, BPM, and SV. However, a clear threshold between the two subgroups was elusive for these features. Surprisingly, left atrial EF failed to distinguish between the two subgroups. Similar trends were noted in the LAA, where lower volumes and SV were observed for stroke cases, yet no distinct threshold emerged for any features, including LAA EF. This suggested the limited utility of EF as a stroke risk assessment biomarker (for both LA and LAA), possibly influenced by the inherent uncertainty associated with its definition.

Turning to hemodynamic simulations, our findings demonstrated that stroke patients generally exhibited lower velocities, increased blood stasis, reduced wall shear stress, and higher values of OSI, RRT, and ECAP. However, this trend applied to all cases except one: an outlier in standalone CFD analysis. Additionally, stroke patients displayed lower kinetic energies, and only the vorticity magnitude at the A-wave could effectively differentiate between the two groups for all patients. Despite a clear pattern observable between stroke and control cases, outliers consistently appeared for almost all features, precluding the identification of some clear-cut hemodynamics features for distinguishing between the two groups, as we summarize in Table 3. To address variability in chamber functionalities, geometries, and contractile properties, we standardized our findings with functional metrics. This approach, characterizing hemodynamic metrics with respect to maximum volume, SV and BPM, provided a clearer distinction between stroke and control cases across all hemodynamic features considered in this study. Similar challenges were noted for both the LA and the LAA in standalone assessments; however, standardizing fluid dynamics results with LAA SV revealed a clear division between the two groups. We provide a clear picture of this standardization in Table 3. Thus, our investigation unveiled a crucial clinical insight: the normalization of hemodynamic features based on ejection fraction failed to differentiate between stroke and control patients. In contrast, normalization with stroke volume revealed a clear and clinically significant distinction, applicable to both the left atrium and its appendage. These findings have noteworthy implications for precise stroke risk assessment in clinical settings.

**Table 3.**
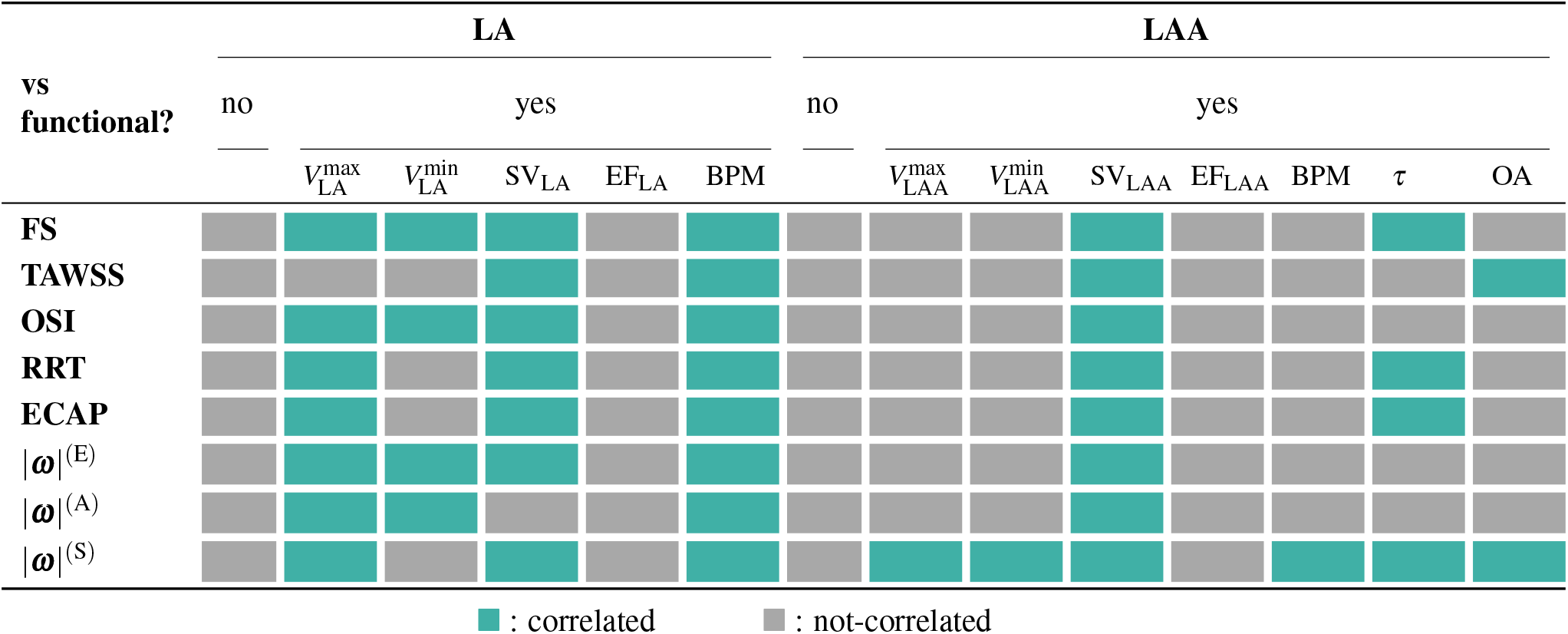
Correlation between hemodynamic features (derived from CFD simulations) and functional measurements (from CMR) for the LA and LAA. 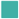 and 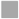 indicate correlation and non-correlation, respectively, in distinguishing stroke by non-stroke cases.

While emphasizing the crucial need to merge these datasets, we underscore the importance of incorporating chamber motion in simulations and data analysis. This highlights the implicit impact of the common assumption of treating the atrium endocardium as rigid in CFD models: a simplification that streamlines the computational process but may lead to potentially inaccurate estimations of stroke risk. Our results align with recent work by Sanatkani et al^37^, which also discusses the potential overestimation of thrombogenic risk in the common assumption of rigid walls.

This work has a few limitations. While incorporating patient-specific atrium motion enhanced the reliability of our results, it constrained us to a relatively small number of patients. Nevertheless, to our knowledge, this study boasts the largest cohort of patients utilizing patient-specific LA motion. As part of future directions, we aim to expand the study to include a larger number of patients, with a consideration for potential sex differences^54,55^. Additionally, we anticipate that utilizing dynamic CT images, as opposed to CINE MRI, might yield different results owing to superior image resolution. However, implementing an Arbitrary Lagrangian Eulerian approach for mesh motion in the CFD model may encounter challenges with intricate geometries and displacements. Our CFD model has its own set of limitations. To streamline the process, we do not explicitly model the MV, which can introduce artifacts in velocity profiles, such as abrupt closures in flowrate profiles. Nevertheless, we believe that this may not significantly impact the computation of hemodynamic features. Moreover, due to a lack of measurements regarding velocity/flow at the pulmonary veins inlets, we make an assumption of a constant physiological pressure for all patients in this paper. This assumption could be relaxed with the availability of pulmonary veins velocity data, for example, from echocardiographic images. Additionally, in terms of data analysis, we posit that further insights could be gleaned from features extracted from Lagrangian simulations, such as the mean age of blood particles – a metric reflecting blood stagnation in the LA and its appendage specifically^21^.

## Methods

We outline the methodologies employed in our analysis. Initially, we detail our CMR data and the devised preprocessing pipeline for the images. Subsequently, we introduce the mathematical models, data assimilation techniques, numerical methods, and simulation software employed for our personalized computational fluid dynamics CFD simulations. Finally, we define key functional and hemodynamic features utilized in assessing stroke risk.

### Patient-specific data and preprocessing

We consider a cohort of 8 patients: 4 control and 4 with history of stroke. These subjects were part of a larger cohort presented in Morris et al.^56^, where they conducted CMR studies using either a 1.5 Tesla or 3 Tesla MR scanner (manufactured by Siemens Healthcare in Erlangen, Germany). The CMR procedures encompassed CINE CMR, CE magnetic resonance angiography (MRA), and 3D late gadolinium enhancement (LGE) CMR. CINE CMR scans were obtained in either the axial or the standard 4-chamber orientation^56^.

These three types of image sequences are combined to eventually produce dynamic meshes. Specifically, a cardiac imaging expert carried out manual segmentation of the LA anatomy from the high-resolution MRA and LGE image. The time-dependent CMR images were then reconstructed from a series of CINE image stacks covering the entire chamber with small or no gap between slices. Then, individual 3D volumes were extracted and reconstructed for each time point of the cardiac cycle from the time-dependent 3D images. Manual segmentation of the LA anatomy from the high-resolution MRA and LGE image was done using the freely-available image processing software itk-SNAP^57,58^. Further details on CMR data and the computational pipeline to recover dynamic meshes can be found in the work by Morris et al^56^. For each patient, we have either 25, 30, 35, or 40 time frames available, depending on the patient. Given the dynamic meshes, we developed a semi-automatic preprocessing pipeline aimed at generating a static volumetric mesh for the CFD simulation and a displacement field (defined with respect to the static mesh configuration) to displace the static mesh over time. In our semi-automatic pipeline, we combine `pvpython`^59^ and `vmtk`^60,61^ preprocessing tools. We generate tetrahedral meshes of the 8 atria with a uniform mesh size approximately equal to 0.8 mm, as we report in Figure 8.

**Figure 8.**
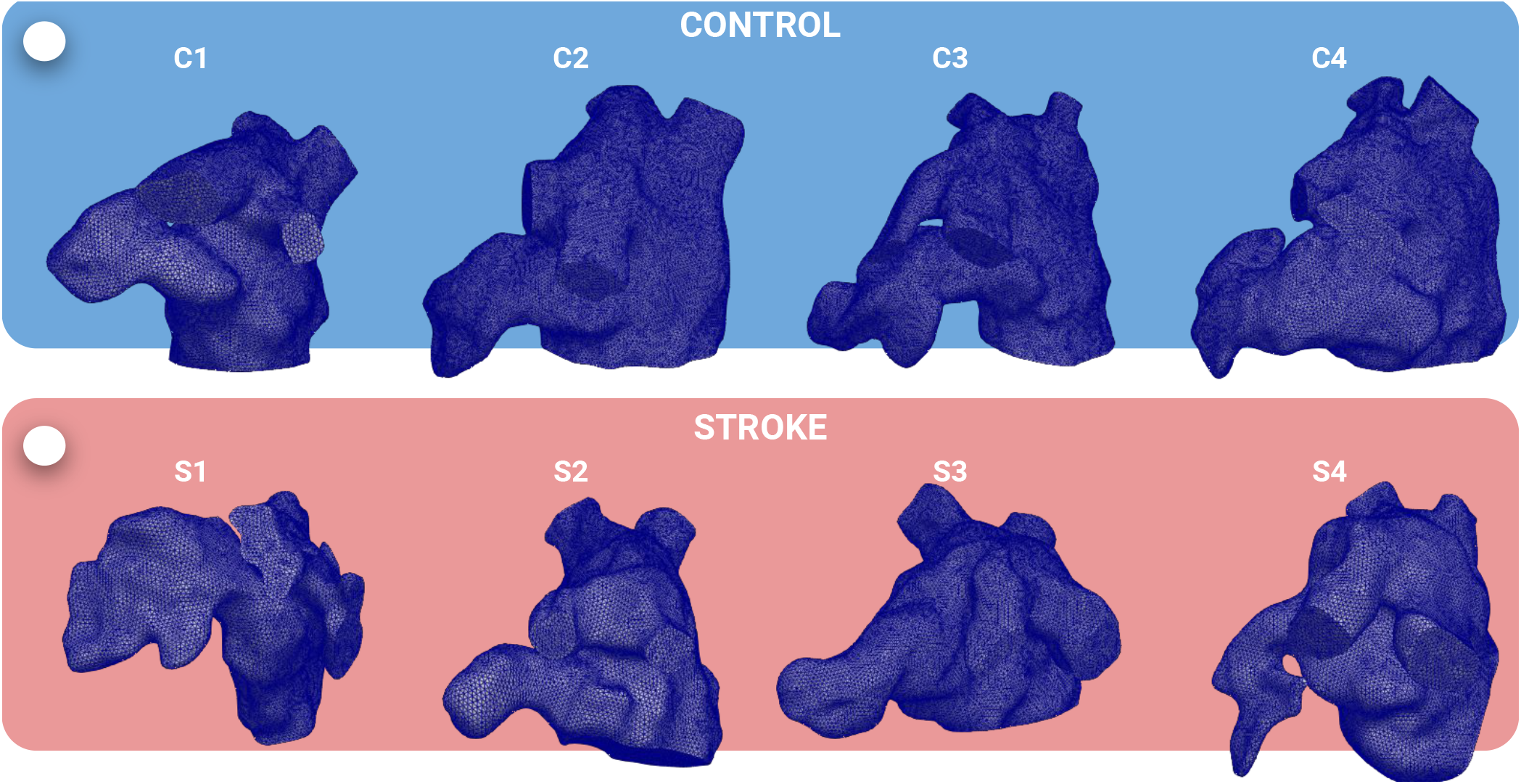
Tetrahedral meshes of the LA for the CFD simulations: control (top) and stroke (bottom) patients.

### Modeling and simulation

We model the blood flow in the LA as an incompressible viscous fluid with constant density *ρ* = 1.06 · 10^3^ kg*/*m^3^ and constant dynamic viscosity *μ* = 3.5 · 10^−3^ kg*/*(ms). We use the incompressible Navier-Stokes equations expressed in the Arbitrary Lagrangian Eulerian (ALE) framework^62^. We set our fluid dynamics domain in Ω_*t*_ delimited by 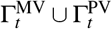. These boundaries are the endocardial wall, MV section, and pulmonary veins (PVs) sections, respectively. The subscript *t* denotes the current (i.e. deformed) configuration of the domain and the subscript 0 denotes the reference (corresponding to the initial) configuration. The domain at any time *t* is defined in terms of a displacement field ***d*** as Ω_*t*_ = {***x*** ∈ ℝ^3^ : ***x*** = ***x***_0_ + ***d***(***x***_0_, *t*), ***x***_0_ ∈ Ω_0_}. We consider a temporal domain (0, *T*_f_), with *T*_f_ = 10*T*_HB_, where *T*_HB_ is the heartbeat period, a patient-specific data. We simulate 10 beats in order to allow the flow to reach a periodic steady-state solution and to neglect the influence of a null velocity initial condition.

The patient-specific displacement is defined for a discrete number of time frames, with a much coarser resolution than the one required by the CFD problem. Thus, we use smoothing splines^63^ to approximate the imaging displacement field in time. We found that this choice – in contrast to other standard approximating or interpolating functions – allows for the reduction of the noise unavoidably introduced in the simulation by the patient-specific data. Let ***d***_*∂*Ω_ be the displacement field on the LA boundary *∂* Ω_*t*_ and on the temporal domain (0, *T*_f_). We compute the displacement of the volumetric mesh ***d*** (i.e. interior mesh points) with the following harmonic lifting problem:

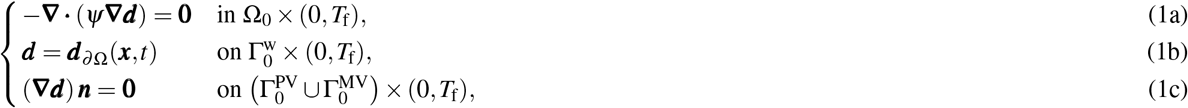

where ***n*** is the outward pointing normal and *ψ*(***x***) is a spatially varying stiffening factor used to avoid mesh element distortion according to the boundary-based stiffening approach proposed in^64^: *ψ*(***x***) = max(*d*(***x***), *α*)^−*β*^ in Ω_0_. *d* is the distance from the LA boundary, *α* and *β* are two parameters that we set equal to *α* = 1.5 mm and *β* = 2, as in^65^. Notice that, in Equation (1), we prescribe natural homogeneous conditions on the PVs and MV sections since the displacement obtained from the images is defined on the endocardial wall only, i.e. where we prescribe Dirichlet conditions. As a matter of fact, 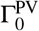 and 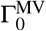 are artificial boundaries required to prescribe inlet and outlet boundary conditions in the CFD problem. The ALE velocity ***u***^ALE^ is then computed by deriving ***d*** with respect to time.

Let ***u*** and *p* be the blood velocity and pressure fields. The incompressible Navier-Stokes equations in the ALE framework endowed with boundary and initial conditions read:

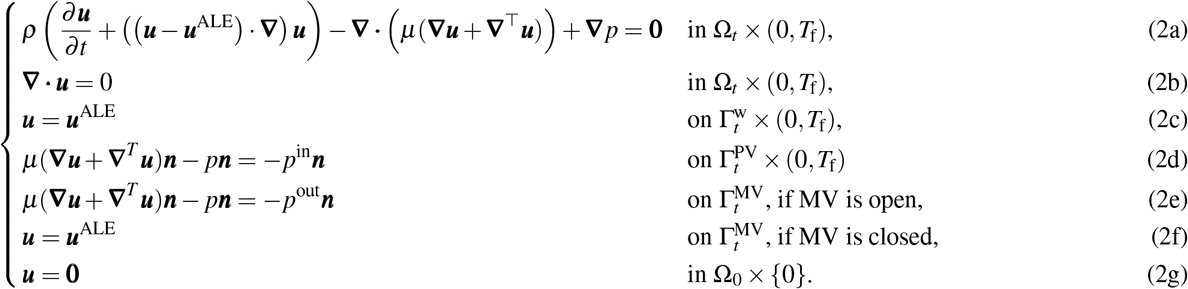

In Figure 9, we provide a graphical sketch of the domain and the boundary conditions we set. We prescribe the ALE velocity on the lateral wall (no-slip condition). On the inlet sections, since we do not have any data in terms of flowrate or pressure, we set Neumann boundary conditions with a pressure *p*^in^ = 10 mmHg, which represents a physiological value measured in the pulmonary veins^66^. On the outlet section, to mimic the opening and closing of the MV, we switch boundary conditions from natural to essential, and vice-versa. Specifically, as done in^50^, we set a no-slip condition if the MV is closed; if the MV is open, we prescribe a physiological value of ventricular pressure during ventricular diastole^67^: *p*^out^ = 5 mmHg. We switch the boundary condition when the LA volume reaches a minimum or maximum, as displayed in Figure 9. Furthermore, we include backflow stabilization in all the Neumann boundaries^68,69^. We start the simulation with a null initial condition.

**Figure 9.**
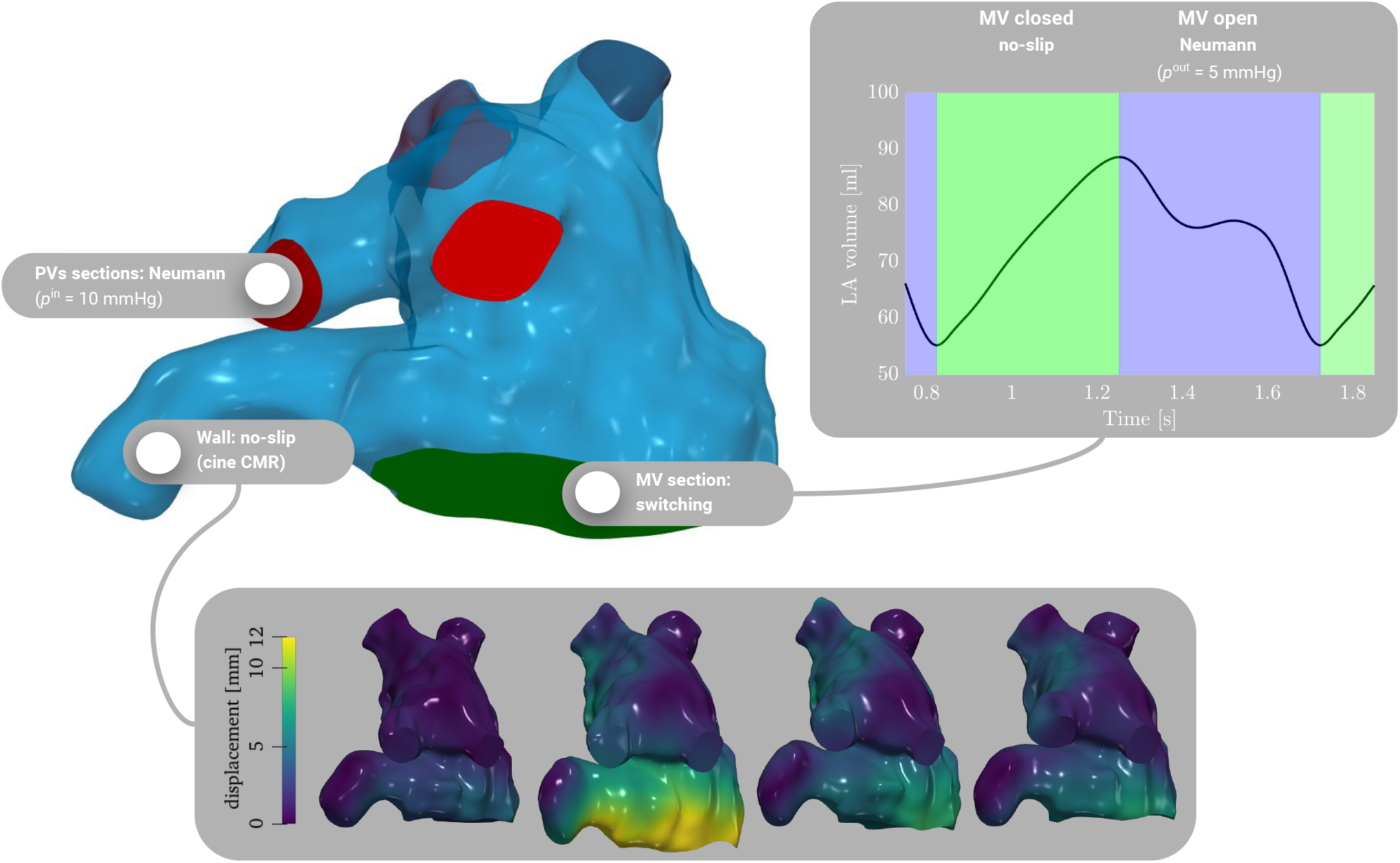
LA computational domain with boundary portions (top). On the inlet section, we prescribe constant pressure equal to 10 mmHg; on the wall, a no-slip condition with patient-specific displacement; on the outlet section, switching boundary condition: no-slip when the MV is closed and pressure equation to 5 mmHg when the MV is open.

We discretize the Navier-Stokes equations in space by the Finite Element (FE) method. We use linear FEs for both velocity and pressure with VMS-LES stabilization^70^, acting also as a turbulence model to account for possible transition-to-turbulence effects in the LA^50^. We treat the nonlinear convection term semi-implicitly, as detailed in^70^. Time integration is carried out with an implicit Euler method, with a constant time step size Δ*t* = 5 · 10^−4^ s.

We solve the CFD problem using `life`^x71^, a high-performance `C++` FE library developed within the iHEART project at Politecnico di Milano. `life`^x^ is mainly focused on cardiac simulations, and based on the `deal.II` finite element core^72–74^. The source code of the life^x^ module for hemodynamics simulations, referred to as `life`^x^`-cfd`, has been recently released^75,76^. Simulation analogous to the one presented in this paper can be carried out with the open source version we released. We run numerical simulations in parallel with 192 cores on the GALILEO100 supercomputer (528 computing nodes each 2 x CPU Intel CascadeLake 8260, with 24 cores each, 2.4 GHz, 384GB RAM) at the CINECA supercomputing center, using 288 cores. The simulation of ten beats for a single patient took about 24 hours.

### Computation of biomarkers

To assess geometrical features of the LAA, we compute its tortuosity as^44^: 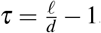, with *ℓ* representing the LAA centerline length and *d* the Euclidean distance of its endpoints. A graphical representation is provided in Figure 10.

**Figure 10.**
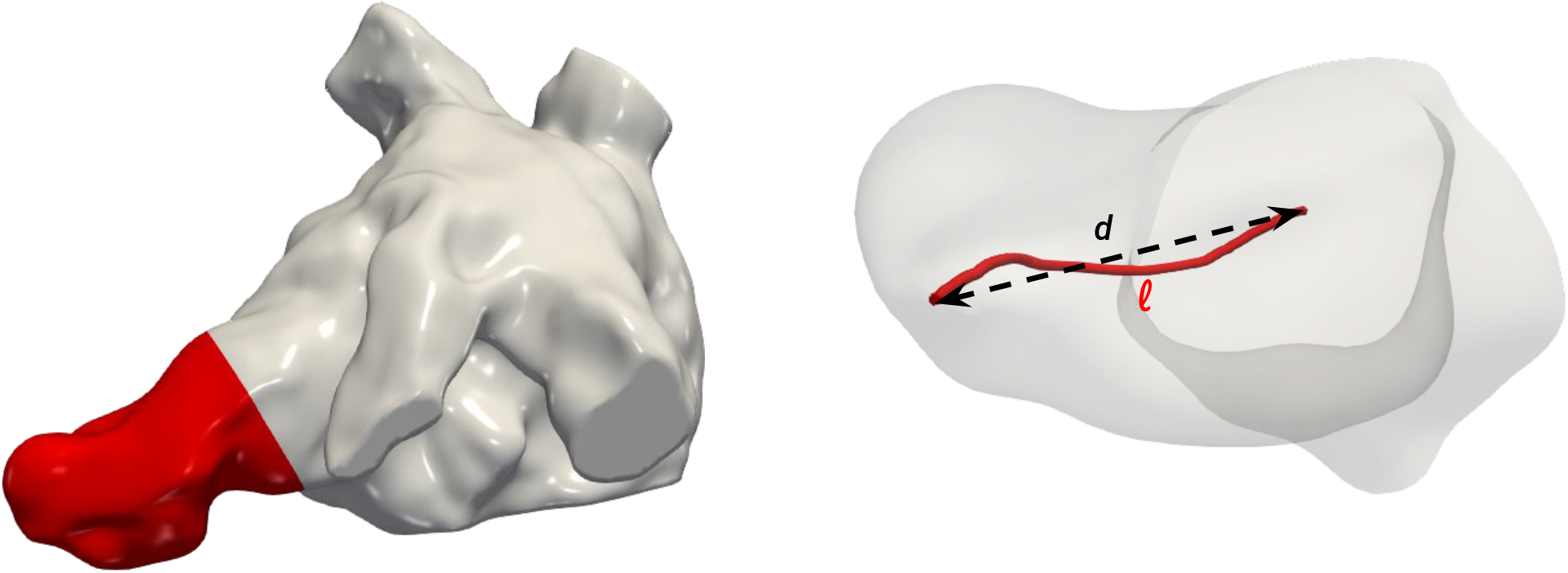
LAA and its centerline for the computation of the tortuosity.

We provide a definition of the hemodynamics features considered in this study. All the features are computed using the *phase-averaged velocity* on the LA boundary in the reference (initial) configuration. We first remove the influence of a null velocity initial condition by discarding the first two heartbeats. Then, we compute the phase-averaged velocity ⟨***u***⟩(***x***, *t*) ∈ Ω_*t*_ ×(0, *T*_HB_) using the remaining eight beats as:

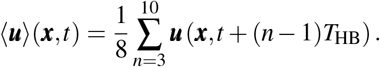

We define the Flow Stasis (FS) as^21^

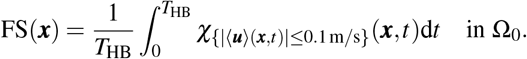

Notice that we defined the flow stasis using a threshold value 0.1 m*/*s, a value consistent with the sensitivity study carried out in^48^. Let *τ*(⟨***u***⟩) = *μ* (**∇**⟨***u***⟩ + **∇**^⊤^⟨***u***⟩) be the viscous stress tensor, the Wall Shear Stress (WSS) vector is defined as

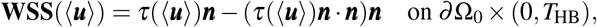

the Time Averaged Wall Shear Stress (TAWSS) as^77^

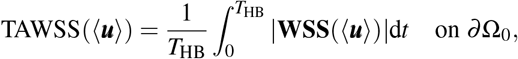

the Oscillatory Shear Index (OSI) as^78^

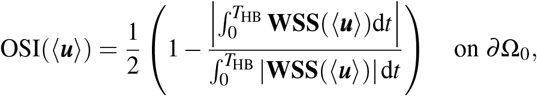

the Relative Residence Time (RRT) as^79^

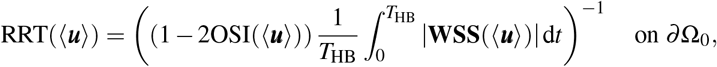

and the Endothelial Cell Activation Potential (ECAP) as^51^

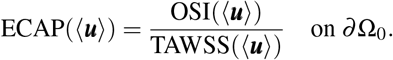

The kinetic energy and the enstrophy are respectively computed as:

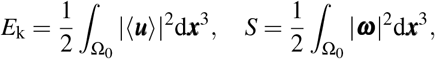

where ***ω*** = ∇ ×⟨***u***⟩ is the phase-averaged vorticity vector.

## Acronyms

ALE: arbitrary lagrangian-eulerian
AF: atrial fibrillation
BPM: beats per minute
CE: contrast enhanced
CFD: computational fluid dynamics
CHA_2_DS_2_-VASc: Congestive Heart Failure, Hypertension, Age ≥ 75 (x2), Diabetes Melitus, Stroke (x2), Vascular Disease, Age (65-74), Sex Category (female +1)
CMR: cardiac magnetic resonance
CPU: central processing unit
CT: computed tomography
EA: early / after wave (ratio)
ECAP: endothelial cell activation potential
EF: ejection fraction
FE: finite element
FS: flow stasis
LA: left atrium
LAA: left atrial appendage
LGE: late gadolinium enhancement
MRA: magnetic resonance angiography
MV: mitral valve
OA: ostium area
OAC: oral anticoagulant
OSI: oscillatory shear index
PV: pulmonary vein
RRT: relative residence time
SV: stroke volume
TAWSS: time-averaged wall shear stress
VMS-LES: variational multiscale large eddy simulation WSS wall shear stress

## Acknowledgements

A.Z., L.D., and A.Q. received funding from the Italian Ministry of University and Research (MIUR) within the PRIN (Research projects of relevant national interest) 2017 “Modeling the heart across the scales: from cardiac cells to the whole organ” Grant Registration number 2017AXL54F.

Z.A. was supported by grant n. T32 HL007024 from the National Heart, Lung, and Blood Institute, National Institutes of Health (NIH) and grant n. T32 GM119998 from the NIH National Institute of General Medicine.

L.D. acknowledges the PRIN 2022 (MIUR Italy) research project “Computational modeling of the human heart: from efficient numerical solvers to cardiac digital twins”; 10.2023–09.2025, Politecnico di Milano, Grant Registration number 202232A8AN. L.D. is member of the INdAM group GNCS “Gruppo Nazionale per il Calcolo Scientifico” (National Group for Scientific Computing), Italy.

N.A.T. acknowledges National Institutes of Health (NIH) grants n. R01HL166759 and R01HL142496 and the Leducq Foundation.

## Author contributions statement

A.Z.: Conceptualization, methodology, software implementation, simulation, formal analysis, writing (original draft). Z.A.: Conceptualization, simulation, formal analysis, writing (original draft). E.K.: Data acquisition and processing, writing (review). K.S.: Data processing, writing (review). L.D.: Conceptualization, supervision, project administration, writing (review). A.Q.: Funding acquisition, conceptualization, supervision, project administration, writing (review). N.T.: Funding acquisition, conceptualization, supervision, project administration, formal analysis, writing (review).

## Additional information

### Accession codes

The datasets used and/or analysed during the current study are available from the corresponding author upon reasonable request. life^x^, the software for carrying out simulations, is freely available at^75,76^.

