## Supplementary Information for "A comprehensive stroke risk assessment by combining atrial computational fluid dynamics simulations and functional patient data"

A. Zingaro, Z. Ahmad, E. Kholmovski, K. Sakata, L. Dede', A. K. Morris, A. Quarteroni, N. A. Trayanova

We report supplementary results from the study. In Supplementary Table 1, we show medians of the boxplot of hemodynamic features computed with our patient-specific CFD simulations in the LA and LAA. In Supplementary Figure 1 we display the boxplot of the hemodynamics features for each patient, focusing on the LAA only (result on the LA are given in the paper). Supplementary Figure 2 shows the LA vorticity magnitude in three different time instants plot against all the functional data considered. Similarly, in Supplementary Figure 3 we show the same results but focused on the LAA (plotted against all the LAA functional data considered).

| | | | **Control** | | | | **Stroke** | | | |
|---|---|---|---|---|---|---|---|---|---|---|
| | | | **C1** | **C2** | **C3** | **C4** | **S1** | **S2** | **S3** | **S4** |
| **LA** | FS | [–] | 0.3019 | 0.3500 | 0.1487 | 0.2960 | 0.4875 | 0.3962 | 0.5254 | 0.3262 |
| | TAWSS | [Pa] | 0.6874 | 0.8157 | 0.8906 | 0.8316 | 0.5922 | 0.5556 | 0.5916 | 0.8205 |
| | OSI | [–] | 0.0804 | 0.0868 | 0.0742 | 0.0636 | 0.0861 | 0.1278 | 0.1371 | 0.0757 |
| | RRT | [1/Pa] | 1.8647 | 1.6056 | 1.4122 | 1.4223 | 2.3328 | 2.6041 | 2.6568 | 1.4575 |
| | ECAP | [1/Pa] | 0.1307 | 0.1048 | 0.0947 | 0.0764 | 0.1673 | 0.2433 | 0.2586 | 0.0899 |
| | $\lvert\boldsymbol{\omega}\rvert^{(E)}$ | [1/s] | 153.2700 | 131.4600 | 141.3100 | 109.1700 | 136.0800 | 118.0900 | 105.8000 | 154.3700 |
| | $\lvert\boldsymbol{\omega}\rvert^{(A)}$ | [1/s] | 107.3800 | 127.7200 | 108.0900 | 124.3200 | 124.6700 | 115.5500 | 133.2500 | 135.3000 |
| | $\lvert\boldsymbol{\omega}\rvert^{(S)}$ | [1/s] | 26.2120 | 23.1450 | 43.6410 | 28.5460 | 18.0620 | 19.7810 | 14.3170 | 30.6820 |
| **LAA** | FS | [–] | 0.3019 | 0.3500 | 1.0000 | 0.8305 | 0.4875 | 0.3962 | 0.7797 | 0.3262 |
| | TAWSS | [Pa] | 0.6874 | 0.8157 | 0.0621 | 0.1110 | 0.5922 | 0.5556 | 0.1770 | 0.8205 |
| | OSI | [–] | 0.0804 | 0.0868 | 0.2130 | 0.0457 | 0.0861 | 0.1278 | 0.0999 | 0.0757 |
| | RRT | [1/Pa] | 1.8647 | 1.6056 | 28.5008 | 11.4786 | 2.3328 | 2.6041 | 8.1796 | 1.4575 |
| | ECAP | [1/Pa] | 0.1307 | 0.1048 | 2.9830 | 0.4140 | 0.1673 | 0.2433 | 0.6936 | 0.0899 |
| | $\lvert\boldsymbol{\omega}\rvert^{(E)}$ | [1/s] | 12.7880 | 16.9670 | 8.7049 | 11.2150 | 6.5866 | 20.7240 | 6.8452 | 6.1957 |
| | $\lvert\boldsymbol{\omega}\rvert^{(E)}$ | [1/s] | 15.0590 | 110.7100 | 10.6680 | 44.7410 | 14.7020 | 85.4180 | 40.5560 | 9.5050 |
| | $\lvert\boldsymbol{\omega}\rvert^{(E)}$ | [1/s] | 14.9870 | 25.4920 | 16.2610 | 11.3740 | 12.2680 | 11.5500 | 10.1500 | 13.9110 |

**Supplementary Table 1.** Medians of boxplot of hemodynamic features computed with patient-specific CFD simulations in the LA and LAA for control and stroke cases.

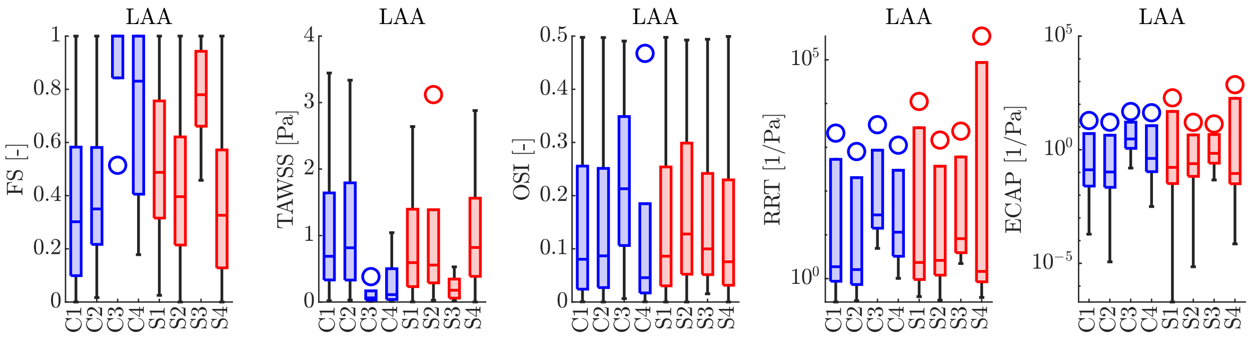


**Supplementary Figure 1.** Boxplots of hemodynamics features computed with patient-specific CFD simulations in the LAA. From the left to the right: FS, TAWSS, OSI, RRT, and ECAP. In blue (■), control cases; in red (■), stroke cases.

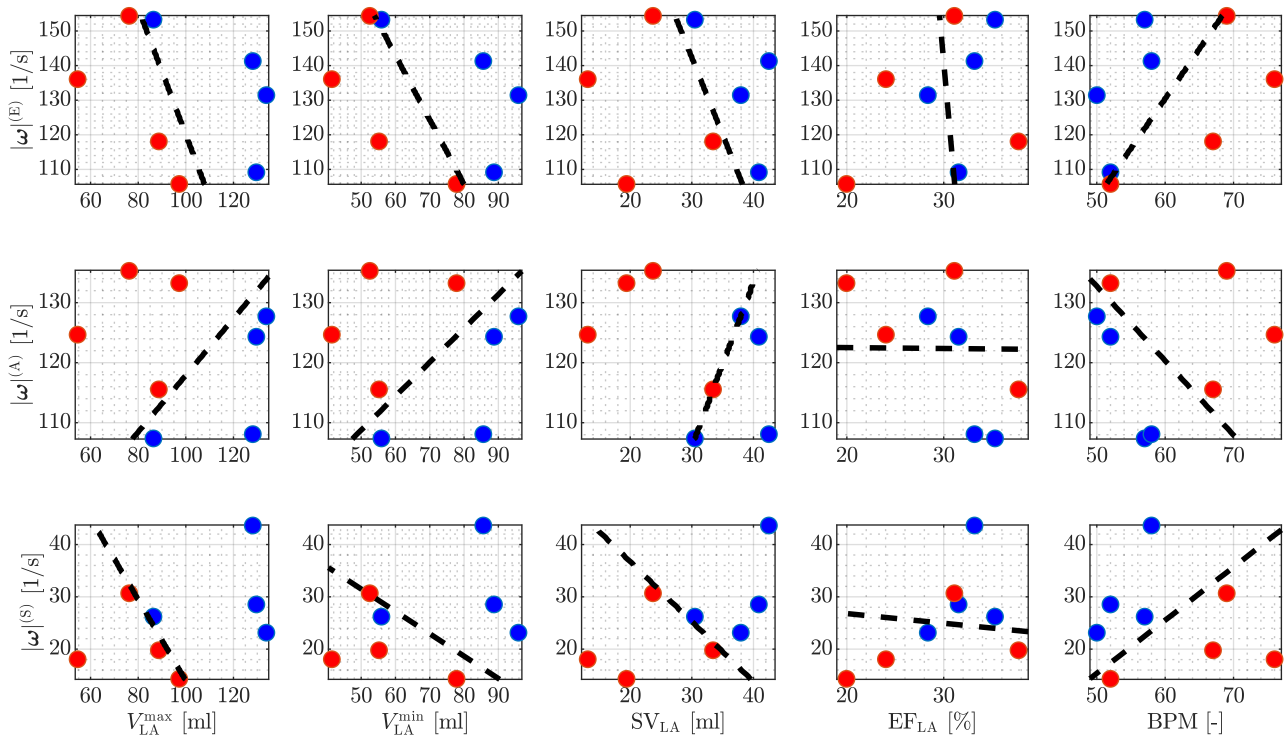


**Supplementary Figure 2.** Medians of vorticity magnitude in the LA against functional data from CMR. In blue (■), control cases; in red (■), stroke cases. The black-dashed line is obtained by running logistic regression. From top to the bottom: vorticity magnitude at E-wave, A-wave, and systolic peak (MV closed) are plotted against maximum LA volume, minimum LA volume, LA SV, LA EF, and BPM (from left to right).

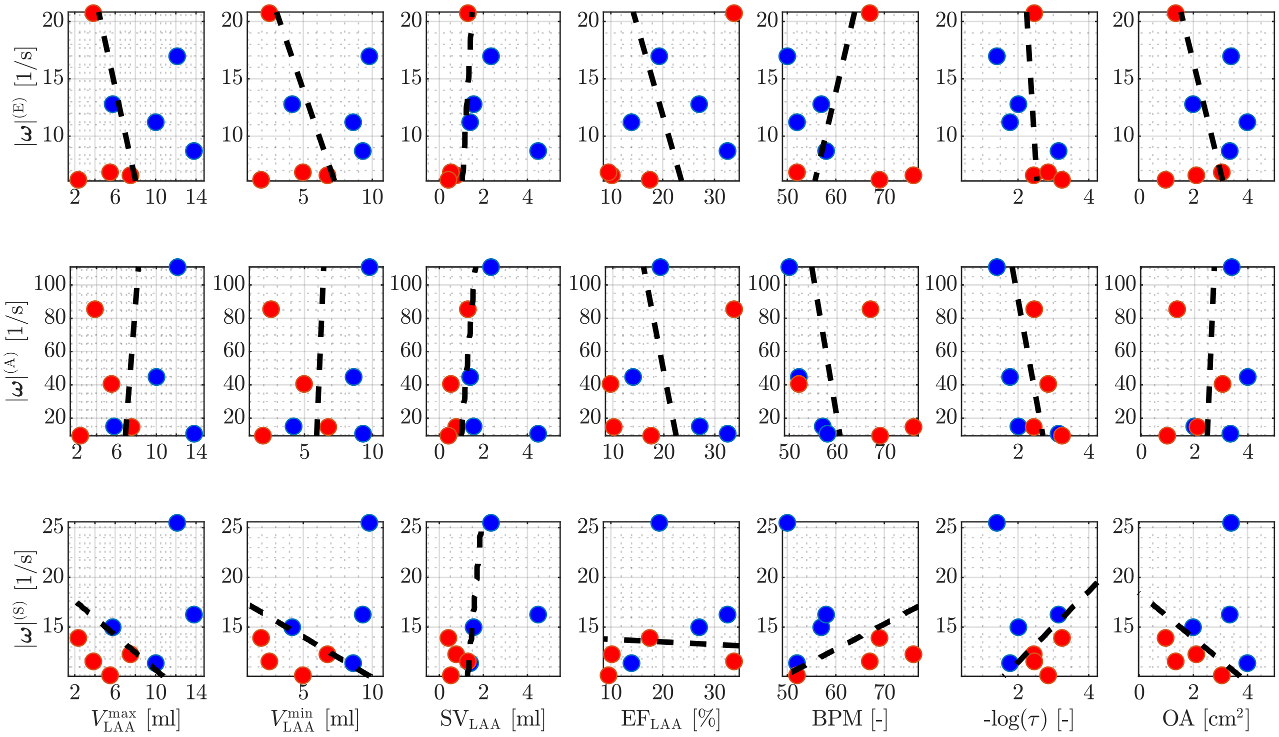


**Supplementary Figure 3.** Medians of vorticity magnitude in the LAA against functional data from CMR. In blue (■), control cases; in red (■), stroke cases. The black-dashed line is obtained by running logistic regression. From top to the bottom: vorticity magnitude at E-wave, A-wave, and systolic peak (MV closed) are plotted against maximum LAA volume, minimum LAA volume, LAA SV, LAA EF, BPM, tortuosity, and OA (from left to right).
